# Synchronicity of viral shedding in molossid bat maternity colonies

**DOI:** 10.1101/2022.08.16.504085

**Authors:** Axel O. G. Hoarau, Marie Köster, Muriel Dietrich, Gildas Le Minter, Léa Joffrin, Riana V. Ramanantsalama, Patrick Mavingui, Camille Lebarbenchon

## Abstract

Infection dynamics in vertebrates are driven by biological and ecological processes. For bats, population structure and reproductive cycles have major effects on RNA virus transmission. On Reunion Island, previous studies have shown that parturition of pregnant females and aggregation of juvenile Reunion free-tailed bats (*Mormopterus francoismoutoui*) are associated to major increase in the prevalence of bats shedding viruses. The synchronicity of such shedding pulses, however, is yet to be assessed, between viruses but also maternity colonies. Based on 3422 fresh faeces collected every two to five weeks during four consecutive birthing seasons, we report the prevalence of bats shedding astroviruses (AstVs), coronaviruses (CoVs), and paramyxoviruses (PMVs) in two maternity colonies on Reunion Island. We found that the proportion of bats shedding viruses is highly influenced by sampling collection dates, and therefore by the seasonality of parturition. We highlight that virus shedding patterns are reproducible among years and colonies for CoVs and at a lesser extent for PMVs, but not for AstVs. We also report 1% of bats harbouring double infections, mostly CoVs and PMVs, but none shedding simultaneously AstVs, CoVs and PMVs.

## INTRODUCTION

Seasonal infection dynamics in vertebrates are driven by a large set of biological and ecological factors, including social behaviours and changes in host population structure [1]. For instance, the breeding season sometimes lead to the aggregation of hundreds to thousands of bats in maternity colonies, generating highly conductive conditions for the transmission of infectious agents [2–6]. Identifying the factors involved in the transmission dynamics of bat-borne pathogens expands our understanding of host-parasite interactions, but is also critical to assess spill-over potential to other hosts [7,8].

Although many studies have reported viruses, bacteria, and blood parasites in bats [9– 11], a precise assessment of the temporal variation in their circulation in host populations remain challenging. In addition to host-related factors, those associated to the bat colony (e.g. type of habitat, occupation length) and to the simultaneous circulation of multiple infectious agents, have not been fully explored. Although co-infections (i.e. presence of at least two infectious agent in a same host) are known to affect host susceptibility or ability to maintain and transmit infectious agents [12–18], their role in infectious agents transmission dynamics in bat populations indeed remains unsolved.

In this study, we investigated the effect of population size, occupation length, and age structure on virus shedding, in molossid bat maternity colonies. Because reproductive cycles are highly synchronized, we hypothesized that viral shedding patterns could be similar and predictable between years. However, differences may be expected between viral families, and associated to either epidemiological or ecological factors such as herd immunity or the maintenance of viral particles in the environment. We also assumed that viral co-circulation in bat populations may generate positive or negative interactions between viruses, with cascade effects on their transmission dynamics.

To test these hypotheses, we focused on the Reunion free-tailed bat (RFTB; *Mormopterus francoismoutoui*), a molossid species endemic to Reunion Island. We collected 3422 fresh faeces simultaneously in two maternity colonies, during four consecutive years, and estimate the prevalence of astroviruses, coronaviruses, and paramyxoviruses. More precisely, (i) we assessed temporal variations in the prevalence of bats shedding viruses, (ii) estimated the proportion of co-infected bats and (iii) tested whether the detection of a given viral family was associated to the presence of another.

## MATERIAL AND METHODS

### Study sites

The study was conducted in two RFTB colonies on Reunion Island, a 2500 km^2^ oceanic island located in the South-Western Indian Ocean. The first colony was located in a ∼ 30 m^3^ natural cave on the West coast of the island (referred to hereafter as “cave”). This colony gather the highest number of bats on the island, and is occupied only during the birthing season, from October to June [6]. During the early stages of the season (October to December), the colony is mainly composed of pregnant females, with a population reaching about 40,000 to 50,000 flying bats before parturition (starting in mid-December) [6]. Then, from January to May, the colony is mainly composed of new-borns and juveniles. Once all juveniles have left the cave, it remains empty until the following maternity season. As for previous studies [6,19], population age structure and parturition timing was visually monitored based on bat morphology (adults: brown fur; new-borns: nude pink skin; juveniles: dark grey fur).

A smaller colony (up to 1,200 flying bats) was also monitored, in a ∼ 5 m^3^ building housing a power transformer, located in the North coast of the island (referred to hereafter as “building”). In this colony, adults are reported all year long but new-borns are present in mid-December, and juveniles between January and May, as for the cave colony. The building remains occupied between June and September, although it is not possible to visually assess the population age structure (differences between juveniles and adults cannot be discriminated based on fur colouration only).

The cave colony was monitored during four consecutive years (i.e. each year corresponding to a birthing season), from October 2016 to March 2020. The building colony was monitored during three consecutive years, from November 2017 to October 2020. Because of the lockdown associated to the COVID19 pandemic, sampling was not done in March and April 2020.

### Sample collection

Fresh bat faeces were collected every two to five weeks, and during the same day in both colonies (except for 9 sampling sessions, because of bad weather limiting bat emergence from the building). For the cave colony, samples were collected in the morning, by placing Benchguard® sheets (60 cm x 49 cm) (Thermo Fisher Scientific Waltham, MA, USA) under roosting bats [19]. For the building colony, samples were collected during bat emergence, at dusk, due to access restriction to the power transformer. Plastic trays with Benchguard® strips (12 cm x 35 cm) were placed below the colony exit. For both colonies, faeces were individually placed into a tube containing 1.5 mL of brain heart infusion medium (Conda, Madrid, Spain) supplemented with penicillin G (1000 units/mL), streptomycin (1 mg/mL), kanamycin (0.5 mg/mL), gentamicin (0.25 mg/mL) and amphotericin B (0.025 mg/mL). Samples were maintained refrigerated in the field and then stored at – 80 °C in the laboratory.

In total, 3422 fresh faeces were collected from the two colonies during 64 sampling sessions (**Supplementary material 1**). In summary, 580 samples were collected in the cave between October 2016 and June 2017 for the first season, during 14 sampling sessions. For the second season, a total of 964 faeces were collected in both sites (cave: n = 499; building: n = 465) between November 2017 and October 2018, during 17 sampling sessions. Nine hundreds and eighty faeces were collected (cave: n = 525; building: n = 455) between November 2018 and September 2019 during 17 sampling sessions. Finally, 898 samples were collected (cave: n = 440; building: n = 458) between October 2019 and October 2020 during 16 sampling sessions.

### RNA extraction and cDNA synthesis

Samples were thawed at 4°C overnight, briefly vortexed, and centrifuged at 1500 g for 15 min. RNA extraction was performed with the IndiSpin QIAcube HT Pathogen Kit as recommended (QIAGEN, Hilden, Germany). Reverse-transcription was performed on 10 µL of RNA, with the ProtoScript II Reverse Transcriptase and Random primers 6 (New England BioLAbs, Ipswich, MA, USA), as previously described [20].

### Virus detection

cDNAs were tested for the presence of the astroviruses (AstVs) RNA-dependent RNA-polymerase (RdRp) gene using a Pan-AstV semi-nested polymerase chain reaction (PCR), following a previously published protocol [21] routinely used in our lab [20,22,23]. PCRs were performed with the GoTaq G2 Hot Start Green Master Mix (Promega, Madison, WI, USA) in an Applied Biosystems 2720 Thermal Cycler (Thermo Fisher Scientific, Waltham, MA, USA). PCR products were visualized on 2% agarose gels stained with 2% Gelred (Biotium, Hayward, CA, USA).

cDNAs were tested for the presence of the coronaviruses (CoVs) RdRp gene using a Pan-Coronavirus multi-probe real-time (RT) PCR following a previously published protocol [24] routinely used in our laboratory [19,25]. PCRs were performed with the QuantiNova Probe PCR Master Mix (QIAGEN, Hilden, Germany) in a CFX96 Touch™ Real-Time PCR Detection System (Bio-Rad, Hercules, CA, USA). Samples collected in the cave between October 2016 and June 2018 were tested for CoV RNA as part of a previous study [19].

cDNAs were tested for the presence of the paramyxoviruses (PMVs) L-polymerase gene targeting *Respiroviruses, Morbilliviruses*, and *Henipaviruses* (RMH) using a semi-nested PCR following a previously published protocol [26] routinely used in our lab [6,27,28], but with slight modifications. A 10-fold dilution of amplicons obtained in the first PCR was used to perform the second PCR and increase specificity. PCRs were performed with the GoTaq G2 Hot Start Green Master Mix (Promega, Madison, WI, USA) in an Applied Biosystems 2720 Thermal Cycler (Thermo Fisher Scientific, Waltham, MA, USA). PCR products were visualized on agarose as above.

### Statistical analysis

Analyses were made with the assumption that each faeces came from an individual bat. Based on the high number of bats (hundreds in the buildings to tens of thousands in the cave), together with the high bat density (> 1,500 individuals per m^2^) [29], and the limited time spent on the colony for sampling (< 15 min), we considered that the probability to collect two or more faeces from the same bat was limited, although this could not be formally excluded. Previous study on CoV shedding dynamics in the cave colony using a similar sampling scheme [19] provided consistent results, supporting the efficiency of such non-invasive methodology to assess the prevalence of bat shedding viruses.

Four generalized linear mixed models (GLMMs) with binomial error structures were used to explore the effect of the colony (different population size and occupation length) respectively on the detection of AstVs, CoVs, and PMVs RNA, as well as co-infections, with the year (i.e. season of parturition: from October to September) and sampling collection dates included as random factors (**Supplementary material 2**). We used three other GLMMs to test if the presence/absence of each virus was influenced by those of the other viruses (**Supplementary material 2**). Analyses were conducted with the package “*lme4*” in R version 4.1.0 [30,31]. The most parsimonious model was selected by excluding, step by step, every variable of the complete model and using an ANOVA with a Chi-square test to compare models (**Supplementary material 2**).

## RESULTS

### Bat population age structure in the colonies

Adult bats were observed between October and December, in both the cave and the building colonies. New-borns were sighted in both colonies from mid-December to late January, suggesting that parturition was highly synchronized between the two maternity colonies (each year, new-borns were sighted for the first time, during the same sampling session in both the cave and the building). From February to May, both colonies were mainly composed of juvenile bats. No bats were observed in the cave between mid-June and September. These observations are consistent with previous studies [6,19].

### Infection dynamics

Astroviruses (AstVs) were detected in both colonies, every year (**Supplementary material 1, Figure 1A**). In total, 57 samples tested positive (mean detection rate ± 95% confidence interval: 1.7% ± 0.4%). A significant variation in AstV RNA detection was found between sampling dates (χ^2^ = 5.23, P < 0.05), with up to 10% ± 10.7% of bats shedding AstV. We did not detect significant variations in AstV RNA detection between years (χ^2^ = 0.08, P = 0.77) nor colonies (χ^2^ = 1.74, P = 0.20).

**Figure 1.**
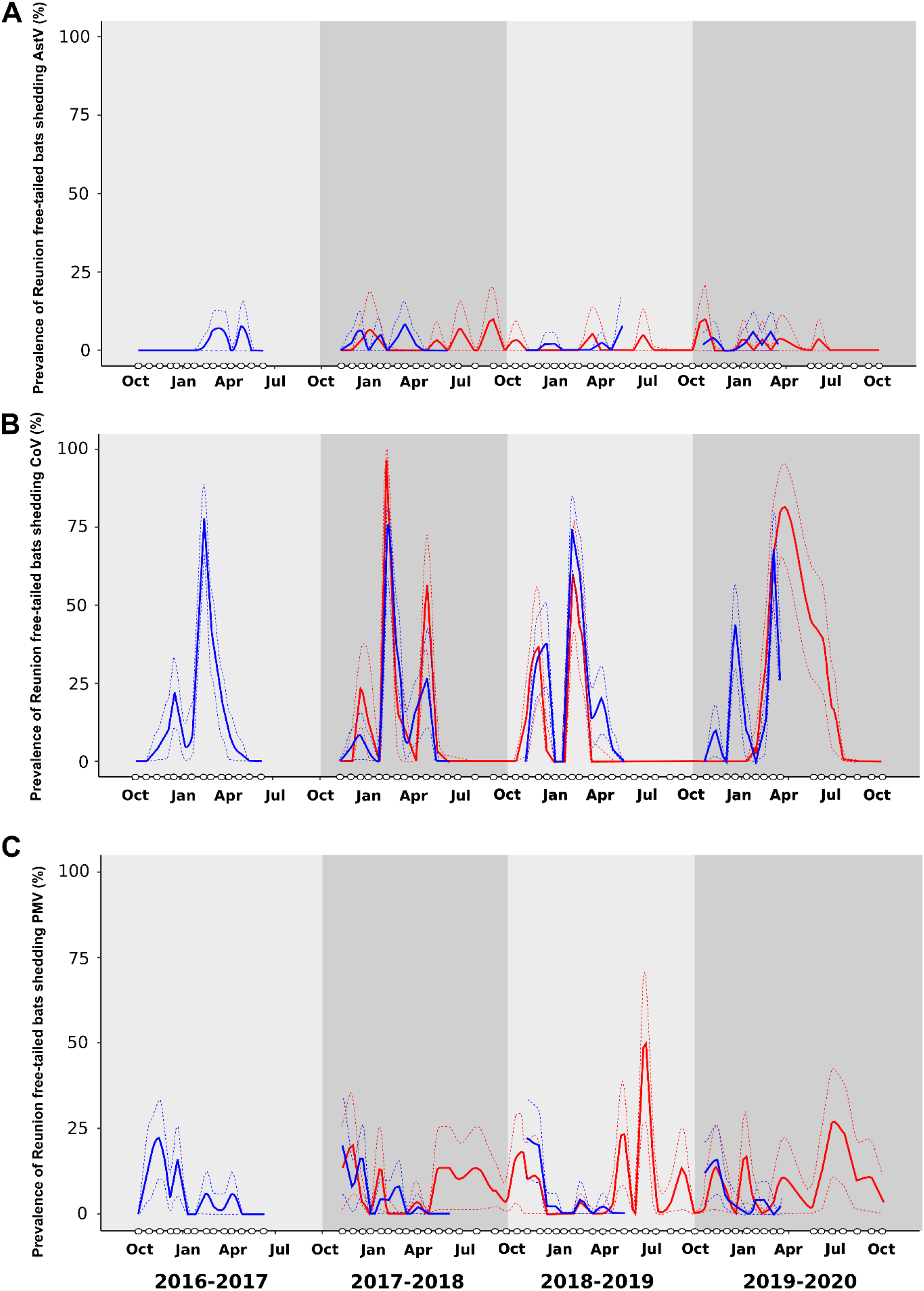
Prevalence of Reunion free-tailed bat (*Mormopterus francoismoutoui*) shedding astrovirus (AstV; A), coronavirus (CoV; B) and paramyxovirus (PMV; C), during four consecutive seasons (October 2016 to October 2020). White dots on X axis indicate sampling dates. Continuous lines represent the proportion of positive samples, and dashed lines the 95% confidence interval. Blue: cave (occupied by bats from October to May); red: building (occupied all year long).

Coronaviruses (CoVs) were detected in both colonies, every year (**Supplementary material 1, Figure 1B**). In total, 573 samples tested positive (16.7% ± 1.3%). A significant variation in CoV RNA detection was found between sampling collection dates (χ^2^ = 975.29, P < 0.001); however, as for AstVs, no significant difference was observed between years (χ^2^ = 0, P = 1) nor colonies (χ^2^ = 0.19, P = 0.66). Major variations in the prevalence of bats shedding CoVs were found, with a first shedding pulse recorded in December (e.g. ranging from 8.0% ± 7.5% to 38.0% ± 13.5% in the cave), followed by a second in February (e.g. ranging from 72.0% ± 12.5% to 78.0% ± 11.5% in the cave). The second shedding pulse was delayed in March, for both colonies, in 2019-2020 (respectively 68.0% ± 13.0% and 80.0% ± 14.3% of bats shedding CoVs in the cave and in the building) (**Supplementary material 1**). None of the samples tested positive between June and October. In the building, the December shedding pulse was not recorded in 2019-2020 (**Figure 1B**).

Paramyxoviruses (PMVs) were detected in both colonies, every year (**Supplementary material 1, Figure 1C**). In total, 224 samples (6.5% ± 0.8%) tested positive. A significant variation in PMV RNA detection was found between sampling collection dates (χ^2^ = 97.69, P < 0.001) but not between years (χ^2^ = 6e-04, P = 0.98). Also, a significant difference was found between colonies (χ^2^ = 5.10, P < 0.05), with 5.4% ± 1.0% of bats shedding PMVs in the cave and 8.3% ± 1.5% in the building. As for CoVs, major variations were detected in the proportion of PMV positive samples within each season (**Figure 1C**). In the cave and the building, a high prevalence was found between November to mid-December (e.g. up to 22.0% ± 11.5% in the cave, and up to 20.0% ± 14.3% in the building). In the building, several other shedding pulses were observed between May-June and September-October (e.g. up to 50.0% ± 21.9% in June in 2018-2019) (**Figure 1C**).

### Co-infections

Co-infections were detected in both colonies and every year. In total, 36 of the 3422 samples tested positive for more than one virus (1.1% ± 0.3%; **Figure 2**). Only bi-infections were observed. Co-infections with CoVs and PMVs (0.7% ± 0.3%) were more common than those with AstVs and PMVs (0.1% ± 0.1%) or AstVs and CoVs (0.3% ± 0.2%) (χ^2^ = 22.19, df = 2, P < 0.001). Overall, a significant variation in co-infection detection was found between sampling collection dates (χ^2^ = 27.48, P <0.001), but not between years (χ^2^ = 0, P = 1), nor colonies (χ^2^ = 0.06, P = 0.81). For the first three years, co-infections were recorded mainly at the beginning of the season in November-December and later in February. In 2019-2020, co-infections were detected in November in the cave, in March for both the cave and the building, and in May and June in the building. There was no statistical association between viruses indicating that the presence of one virus may not be determined by the presence of another (**Supplementary material 3**).

**Figure 2.**
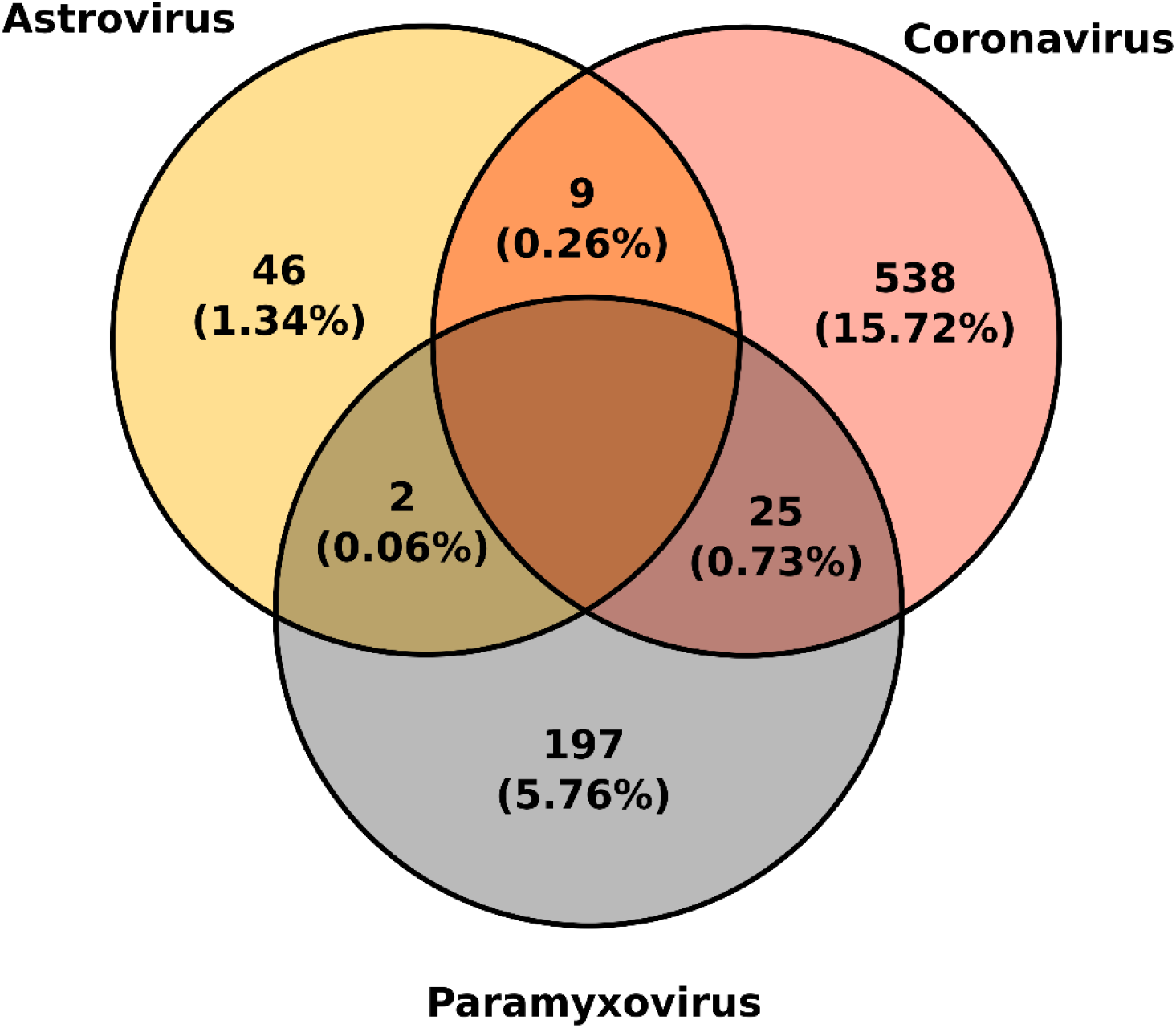
Co-infections between astroviruses, coronaviruses and paramyxoviruses in Reunion freetailed bat (*Mormopterus francoismoutoui*). The number and proportion of positive samples is indicated for each virus.

## DISCUSSION

Based on the molecular detection of AstVs, CoVs and PMVs, we precisely assessed temporal variations in virus shedding dynamics in two RFTB colonies during four consecutive years. In spite of differences between colonies, notably population size (up to 50,000 *vs* 1,200 flying bats), and occupation length (maternity period vs permanent colony), a similar shedding pattern was observed between colonies for CoVs and, at a lesser extent, for PMVs. Temporal variation was likely associated to the high synchrony of reproductive cycles and parturition of the studied bat species, although differences between viral families could be driven by host, viral or environmental factors. We also detected a slight proportion of co-infected bats, with CoVs and PMVs co-infections more commonly detected than AstVs and CoVs, or AstVs and PMVs.

A major shedding pulse was observed almost each year, in February, for CoVs. This pattern was highly repeatable between years, but also synchronized between colonies. Population size may therefore not affect CoVs transmission in RFTB, but rather, age structure could be the main driver of epidemiological cycles, as previously suggested [19]. This major shedding pulse could indeed be associated to the waning of maternal antibodies in juvenile bats, although the duration of maternal antibodies of bat CoVs is yet to be assessed. Interestingly, in 2019-2020, this major shedding pulse was slightly delayed, in March, in both colonies. This may reflect a late arrival of pregnant females, resulting in delayed births and waning of maternal antibodies in juvenile bats. It may also be explained by the introduction of naive juvenile bats arriving from other colonies, prospecting these sites as new potential roosts. Shedding pulses were also detected earlier in the season, at the time of parturition, likely corresponding to CoV infection in pregnant females [19]. This pulse was also repeatable between years and synchronized between colonies, although not detected in the building in 2019-2020. This might also reflect a late arrival of pregnant females in the colony, corroborating the slightly delayed shedding pulse detected in juvenile bats, later in the season.

Repeated shedding pulses were also detected for PMVs, although less pronounced than for CoVs. Synchronized increases were observed between colonies, at the beginning of the birthing season (November) and could be associated to the arrival of pregnant females [6], as for CoVs. Although more inconsistently, PMVs were also detected after parturition, between January and May. These increases were minor as compared to CoVs, but may also be associated to the waning of maternal antibodies in juvenile bats [6].

In addition to the variations in the prevalence of bats shedding PMVs and CoVs, we also highlighted a major difference after the birthing season, between June and October. Indeed, PMVs were detected in the building, every year, but not CoVs, raising questions about the maintenance of viral transmission in bat populations for each of these viruses. Two different mechanisms may indeed be involved: a maintenance and transmission of PMVs in roosting habitats mostly composed by juvenile bats, and an environmental persistence of CoV viral particles, leading to yearly exposure of susceptible individuals at the beginning of the birthing season. These hypotheses could be tested by developing further field studies focusing on bat population structure as well as on experimental studies assessing the role of abiotic factors in virus maintenance outside the host body.

For AstVs, despite changes in the prevalence of bats shedding viruses, no clear pattern associated to variation in the population age structure, nor colony type, was evidenced. These results contrast with a previous longitudinal study conducted in Germany, in a Mouse-eared bat (*Myotis myotis*) maternity colony, reporting AstV shedding pulses early in the birthing season [4]. These differences could be related to variations in host susceptibility, but also be associated to environmental conditions influencing the persistence of non-enveloped viral particles, such as AstVs. Colony type (size, air flow, humidity, temperature, etc.) could affect environmental persistence and transmission opportunities for these viruses, in particular between tropical and temperate regions. A larger set of bat colonies could be investigated, with measurements of environmental variables (e.g. temperature, humidity, etc.) to assess the role of these factors in AstV transmission.

Co-infections involving CoVs and PMVs were mostly detected, and may be explained by the higher proportions of bats shedding these two viruses, as compared to AstVs. We did not report any interaction among viruses. In Borneo, a study conducted in eight tropical species reported co-infections and interactions between AstVs and CoVs in Fawn Leaf-nosed bats (*Hipposideros cervinus*) [13]. We focused on the co-circulation of three viral families, but bats are natural hosts of a large diversity of infectious agents, potentially interacting [9,32]. The RFTB has for example been found to host *Leptospira* sp. bacteria whose presence is positively associated to the presence of PMVs [6]. Experimental approaches without a priori (*i*.*e*. next generation sequencing, metagenomics) may provide the opportunity to better characterize the range of infectious agents that may be involved in co-infections [15,16], and their consequences on transmission dynamics.

Overall, this study shows a strong link between the seasonality of parturition and virus infection dynamics in a tropical bat population. Different shedding patterns between viral families were found, with predictable infection patterns among years for CoVs. These findings also underline the need to better assess the drivers involved in virus transmission dynamics in bats, beyond host-related factors, by focusing on the interactions with the environment and other infectious agents. Such a global assessment is needed to explore spill-over risk associated to major shedding pulses in bat populations [7,16].

## ACKNOWLEDGMENTS

We thank Thomas Juhasz, Gaëlle Lefèvre, Maëlle Teysseire and Céline Toty for their assistance in the field or in the laboratory.

## ETHICAL STATEMENT

This study is based on environmental sampling only, and therefore not subjected to approval by an ethic committee.

## DATA ACCESIBILITY

Data are available in the supplementary material.

## AUTHORS’ CONTRIBUTION

AOGH, CL and PM conceived the study. AOGH, CL, GLM, LJ, MD and RVR collected biological material in the field. AOGH, CL, LJ, MK and RVR performed the molecular analyses. AOGH, CL and MD analysed the data and drafted the manuscript. All authors edited, read and approved the final version of the manuscript.

