## Supplementary material 1 for "Synchronicity of viral shedding in molossid bat maternity colonies"

**Supplementary material 1.** Number of tested (N_tested_) and positive samples (N_positive_) (AstV, CoV and PMV) for each sampling session, in the cave and building colonies. * dates for which the sampling collection couldn’t be done during the same day in both colonies.

|  |  |  |  |  | **Npositive** | | | | | |
| --- | --- | --- | --- | --- | --- | --- | --- | --- | --- | --- |
|  |  | **Ntested** | |  | **Cave** | | | **Building** | | |
|  | **Date of collection** | **Cave** | **Building** | **All sites** | **AstV** | **CoV** | **PmV** | **AstV** | **CoV** | **PmV** |
| **2016-2017** | 6-Oct-16 | 12 | N/A | 12 | 0 | 0 | 0 | N/A | N/A | N/A |
|  | 27-Oct-16 | 39 | N/A | 39 | 0 | 0 | 6 | N/A | N/A | N/A |
|  | 17-Nov-16 | 50 | N/A | 50 | 0 | 2 | 11 | N/A | N/A | N/A |
|  | 8-Dec-16 | 40 | N/A | 40 | 0 | 4 | 2 | N/A | N/A | N/A |
|  | 20-Dec-16 | 50 | N/A | 50 | 0 | 11 | 8 | N/A | N/A | N/A |
|  | 10-Jan-17 | 40 | N/A | 40 | 0 | 2 | 0 | N/A | N/A | N/A |
|  | 24-Jan-17 | 50 | N/A | 50 | 0 | 5 | 0 | N/A | N/A | N/A |
|  | 16-Feb-17 | 50 | N/A | 50 | 1 | 39 | 3 | N/A | N/A | N/A |
|  | 2-Mar-17 | 49 | N/A | 49 | 3 | 20 | 1 | N/A | N/A | N/A |
|  | 23-Mar-17 | 50 | N/A | 50 | 3 | 9 | 1 | N/A | N/A | N/A |
|  | 6-Apr-17 | 50 | N/A | 50 | 0 | 4 | 3 | N/A | N/A | N/A |
|  | 26-Apr-17 | 40 | N/A | 40 | 3 | 1 | 0 | N/A | N/A | N/A |
|  | 16-May-17 | 35 | N/A | 35 | 0 | 0 | 0 | N/A | N/A | N/A |
|  | 8-Jun-17 | 25 | N/A | 25 | 0 | 0 | 0 | N/A | N/A | N/A |
| **Total** | **14 sampling sessions** | **580** | **N/A** | **580** | **10** | **97** | **35** | **N/A** | **N/A** | **N/A** |
| **2017-2018** | 9-Nov-17 | 30 | 30 | 60 | 0 | 0 | 6 | 0 | 0 | 4 |
|  | 30-Nov-17 | 50 | 30 | 80 | 1 | 2 | 4 | 0 | 0 | 6 |
|  | 19-Dec-17 | 50 | 30 | 80 | 3 | 4 | 8 | 1 | 7 | 1 |
|  | *2-Jan-18 | N/A | 15 | 49 | N/A | N/A | N/A | 1 | 2 | 0 |
|  | *3-Jan-18 | 34 | N/A |  | 0 | 1 | 0 | N/A | N/A | N/A |
|  | 24-Jan-18 | 45 | 30 | 75 | 2 | 0 | 2 | 1 | 0 | 4 |
|  | 7-Feb-18 | 50 | 30 | 80 | 0 | 36 | 2 | 0 | 29 | 0 |
|  | 28-Feb-18 | 50 | 20 | 70 | 2 | 18 | 4 | 0 | 3 | 0 |
|  | 14-Mar-18 | 50 | 10 | 60 | 3 | 4 | 0 | 0 | 1 | 0 |
|  | 5-Apr-18 | 50 | 30 | 80 | 1 | 8 | 1 | 0 | 0 | 1 |
|  | 26-Apr-18 | 30 | 30 | 60 | 0 | 8 | 0 | 0 | 17 | 0 |
|  | 17-May-18 | 30 | 30 | 60 | 0 | 0 | 0 | 1 | 1 | 4 |
|  | 6-Jun-18 | 30 | 30 | 60 | 0 | 0 | 0 | 0 | 0 | 4 |
|  | 26-Jun-18 | N/A | 30 | 30 | N/A | N/A | N/A | 2 | 0 | 3 |
|  | 30-Jul-18 | N/A | 30 | 30 | N/A | N/A | N/A | 0 | 0 | 4 |
|  | 4-Sep-18 | N/A | 30 | 30 | N/A | N/A | N/A | 3 | 0 | 2 |
|  | 26-Sep-18 | N/A | 30 | 30 | N/A | N/A | N/A | 0 | 0 | 1 |
|  | 17-Oct-18 | N/A | 30 | 30 | N/A | N/A | N/A | 1 | 0 | 5 |
| **Total** | **17 sampling sessions** | **499** | **465** | **964** | **12** | **81** | **27** | **10** | **60** | **39** |
| **2018-2019** | 6-Nov-18 | 50 | 30 | 80 | 0 | 0 | 11 | 0 | 4 | 3 |
|  | *30-Nov-18 | 50 | N/A | 80 | 0 | 17 | 10 | N/A | N/A | N/A |
|  | *1-Dec-18 | N/A | 30 |  | N/A | N/A | N/A | 0 | 11 | 3 |
|  | 17-Dec-18 | 50 | 30 | 80 | 0 | 19 | 1 | 0 | 1 | 0 |
|  | 2-Jan-19 | 50 | 30 | 80 | 1 | 0 | 1 | 0 | 0 | 0 |
|  | 17-Jan-19 | 50 | 30 | 80 | 0 | 0 | 0 | 0 | 0 | 0 |
|  | *4-Feb-19 | 50 | N/A | 80 | 0 | 37 | 0 | N/A | N/A | N/A |
|  | *5-Feb-19 | N/A | 30 |  | N/A | N/A | N/A | 0 | 18 | 0 |
|  | 18-Feb-19 | 50 | 30 | 80 | 0 | 30 | 2 | 0 | 13 | 1 |
|  | 14-Mar-19 | 50 | 20 | 70 | 0 | 7 | 0 | 1 | 0 | 0 |
|  | *2-Apr-19 | 50 | N/A | 55 | 1 | 10 | 1 | N/A | N/A | N/A |
|  | *3-Apr-19 | N/A | 5 |  | N/A | N/A | N/A | 0 | 0 | 0 |
|  | 23-Apr-19 | 50 | 30 | 80 | 0 | 2 | 0 | 0 | 0 | 1 |
|  | 15-May-19 | 25 | 30 | 55 | 2 | 0 | 0 | 0 | 0 | 7 |
|  | 5-Jun-19 | N/A | 30 | 30 | N/A | N/A | N/A | 0 | 0 | 0 |
|  | 26-Jun-19 | N/A | 20 | 20 | N/A | N/A | N/A | 1 | 0 | 10 |
|  | 16-Jul-19 | N/A | 20 | 20 | N/A | N/A | N/A | 0 | 0 | 1 |
|  | 14-Aug-19 | N/A | 30 | 30 | N/A | N/A | N/A | 0 | 0 | 1 |
|  | 5-Sep-19 | N/A | 30 | 30 | N/A | N/A | N/A | 0 | 0 | 4 |
|  | 30-Sep-19 | N/A | 30 | 30 | N/A | N/A | N/A | 0 | 0 | 0 |
| **Total** | **17 sampling sessions** | **525** | **455** | **980** | **4** | **122** | **26** | **2** | **47** | **31** |
| **2019-2020** | *21-Oct-19 | 50 | N/A | 80 | 1 | 0 | 6 | N/A | N/A | N/A |
|  | *24-Oct-19 | N/A | 30 |  | N/A | N/A | N/A | 3 | 0 | 1 |
|  | 12-Nov-19 | 50 | 30 | 80 | 2 | 5 | 8 | 0 | 0 | 4 |
|  | *2-Dec-19 | 50 | N/A | 80 | 0 | 0 | 2 | N/A | N/A | N/A |
|  | *3-Dec-19 | N/A | 30 |  | N/A | N/A | N/A | 0 | 0 | 2 |
|  | *19-Dec-19 | 50 | N/A | 57 | 0 | 22 | 1 | N/A | N/A | N/A |
|  | *20-Dec-19 | N/A | 7 |  | N/A | N/A | N/A | 0 | 0 | 0 |
|  | 10-Jan-20 | 40 | 30 | 70 | 1 | 4 | 0 | 1 | 0 | 5 |
|  | *27-Jan-20 | N/A | 30 | 80 | N/A | N/A | N/A | 0 | 1 | 1 |
|  | *28-Jan-20 | 50 | N/A |  | 3 | 0 | 2 | N/A | N/A | N/A |
|  | 13-Feb-20 | 50 | 30 | 80 | 1 | 5 | 2 | 1 | 8 | 0 |
|  | *2-Mar-20 | N/A | 30 | 80 | N/A | N/A | N/A | 0 | 20 | 1 |
|  | *3-Mar-20 | 50 | N/A |  | 3 | 34 | 0 | N/A | N/A | N/A |
|  | 16-Mar-20 | 50 | 30 | 80 | 1 | 13 | 1 | 1 | 24 | 3 |
|  | 20-May-20 | N/A | 31 | 31 | N/A | N/A | N/A | 0 | 13 | 1 |
|  | 4-Jun-20 | N/A | 30 | 30 | N/A | N/A | N/A | 1 | 12 | 4 |
|  | 25-Jun-20 | N/A | 30 | 30 | N/A | N/A | N/A | 0 | 5 | 8 |
|  | 16-Jul-20 | N/A | 30 | 30 | N/A | N/A | N/A | 0 | 0 | 7 |
|  | 13-Aug-20 | N/A | 30 | 30 | N/A | N/A | N/A | 0 | 0 | 3 |
|  | 10-Sep-20 | N/A | 30 | 30 | N/A | N/A | N/A | 0 | 0 | 3 |
|  | 1-Oct-20 | N/A | 30 | 30 | N/A | N/A | N/A | 0 | 0 | 1 |
| **Total** | **16 sampling sessions** | **440** | **458** | **898** | **12** | **83** | **22** | **7** | **83** | **44** |
| **Total** | **64 sampling sessions** | **2044** | **1378** | **3422** | **38** | **383** | **110** | **19** | **190** | **114** |
