## Supplementary material 2 for "Synchronicity of viral shedding in molossid bat maternity colonies"

**Supplementary material 2.** Generalized linear mixed models summaries. AstV: astrovirus; CoV: coronavirus; PMV: paramyxovirus.

| **Response variable** | **Fixed effect** | **Random effect** |  | **Complete model** | **Selected model** |
| --- | --- | --- | --- | --- | --- |
| AstV | Colony | (Year/Date) |  | AstV ~ Colony+(1\|Year/Date) | AstV ~ 1+(1\|Date) |
| CoV | Colony | (Year/Date) |  | CoV ~ Colony +(1\|Year/Date) | CoV ~ 1+(1\|Date) |
| PMV | Colony | (Year/Date) |  | PMV ~ Colony +(1\|Year/Date) | PMV ~ Colony +(1\|Date) |
| Coinfection | Colony | (Year/Date) |  | Coinfection ~ Colony +(1\|Year/Date) | Coinfection ~ 1+(1\|Date) |
| AstV | CoV+PMV+CoV:PMV | (Date) |  | AstV ~ CoV+PMV+CoV:PMV+(1\|Date) | AstV ~ 1+(1\|Date) |
| CoV | AstV+PMV+AstV:PMV | (Date) |  | CoV ~ AstV+PMV+AstV:PMV+(1\|Date) | CoV ~ 1+(1\|Date) |
| PMV | AstV+CoV+AstV:CoV | (Date) |  | PMV ~ AstV+CoV+AstV:CoV+(1\|Date) | PMV ~ 1+(1\|Date) |
