## Supplementary material 3 for "Synchronicity of viral shedding in molossid bat maternity colonies"

**Supplementary material 3.** Statistics of generalized linear models testing for associations between astroviruses, coronaviruses, and paramyxoviruses.

| **Type of association** | **Chi-square value (χ²)** | **P-value** |
| --- | --- | --- |
| AstV-CoV | 0.06 | 0.81 |
| AstV-PMV | 1.04 | 0.31 |
| AstV-CoV:PMV | 0.46 | 0.5 |
| CoV-AstV | 0.25 | 0.62 |
| CoV-PMV | 0.02 | 0.9 |
| CoV - AstV:PMV | 0.51 | 0.47 |
| PMV-AstV | 560.38 | 0 |
| PMV-CoV | 0.43 | 0.51 |
| PMV-AstV:CoV | 0.58 | 0.45 |
